# Eye movement based information system indicates human behavior in virtual driving

**DOI:** 10.1101/2022.07.18.498964

**Authors:** Zhe Peng, Qing Xu, Runlin Zhang, Klaus Schoeffmann, Simon Parkinson

**Affiliations:** Tianjin University; University of Huddersfield; Alpen-Adria-Universitaet Klagenfurt

## Abstract

Humans modulate the behavior flexibly after timely receiving and processing information from the environment. To better understand and measure human behavior in the driving process, we integrate humans and the environment as a system. The eye-movement methodologies are used to provide a bridge between humans and environment. Thus, we conduct a goal-directed task in virtual driving to investigate the law of eye-movement that could characterize the humans (internal) and environmental (external) state measured by fixation distribution and optical flows distribution. The analysis of eye-movement data combined with the information-theoretic tool, transfer entropy, active information storage, quantify the humans’ cognitive effort and receiving information, and in fact, there is a balance (optimal) range between two, because of the mutual synergy and inhibition, whose quantified value is named balance of information processing. Subsequently, we update a system-level model, finding that those information measurements, transfer entropy, active information storage, and balance of information processing, all are included. This information set is information flow, which is quantified by the square root of Jensen-Shannon divergence (SRJSD), named information flow gain. What’s more, results also demonstrate that the influence of system-level information flow correlated with behavioral performance stronger than the separate measurements. In conclusion, we research humans’ eye-movement based on information theory to analyze behavioral performance. Besides driving, these measurements may be a predictor for other behaviors such as walking, running, etc. Still, the limitation is that the information flow may be a proxy of determinants of behavior.

## Introduction

The human behavior is complex implying to a cognitive process such as driving process, where drivers continuously acquire information from surrounding environment [1, 2]. Eye-movement could characterize the degree of load, attentions, awareness, perception in driving processes [3]. Indeed, it reflects humans’ cognition in real world [4]. Whereas we will meet unpredictable conditions such as traffic jams or crowds in driving process, which make the eye-movement pattern so complicated that include too much information irrelevant to driving. At present, the virtual reality could eliminate the other distractors and only supply the stimuli that are applied to the task for its controllable properties [5]. It also could construct the immersive 3D scene approaching more realistic reaction of humans in real-time because the 3D objects allow in-scene depth, multiple angles, and variation of material’s properties [6]. What’s more, estimating gaze position in 3D scene coordinates used eye-tracking device is more accurate [7].

The goal of this study is to examine the task performance by analyzing the eye-movement strategies. Detailed, previous literature shows how different eye-movement strategies that could be appropriate for different stimulus formats that finally affect task performance [8], and eye-movement combined with information theory based on Shannon entropy could indicate the effects such as prediction, error reduction, and decision-making relating to task performance [9–11]. For examples, as predictors of driving performance, the stationary gaze entropy (SGE) and gaze transition entropy (GTE) are used to improve the regression and classification model in alcohol and sleep conditions [12, 13]. Not only in driving tasks, while craft-workers and doctors were performing their specific tasks, whose eye-movement strategies is also tightly relevant to the goal that to be achieved [14, 15]. So, it is meaningful to explore the law of cognitive process of these specific skills in future work.

For those cases, the interaction of humans and environment constitutes a complex information system, not separated. On the one hand, the different kinds or amounts of environmental information may change the humans’ state such as cognitive load that even influence the ability of information processing (modulation control) [16]. On the other hand, humans actively control their behavior, such as oculomotor or motor action, which could change the observed environmental information quantified by optical flow on retina [17]. Moreover, the active information storage (AIS) and transfer entropy (TE) are the information-theoretic tools widely used to quantify the information storage and transfer in one or two stochastic processes in this complex system [18–22]. However, there is not lots of studies about information processing. They pay more attention to the behavioral changes after humans have received and processed information [23, 24].Thus especially, we propose a novel metric for information processing, where one of the factors is the control setting that a pattern of eye-movement strategies could characterize [25]. For instance, the humans prefer to adjust the fixation oriented toward the location with multiple relevant or effective information. In driving process, the driver first considers the riskiest location and then fixation on it to ensure safety. During this time, humans are also affected by cognitive effort that also could be characterized by eye-movement pattern as another factor. Any extreme extent of effort will interfere the ability of control, in other words, there are an optimal range to coordinate the relation between control setting and cognitive effort [9]. To this aim, the research of [24] confirms a balanced mode could ensure the foot-placement accuracy based on the effort of body movement and eye-movement strategies, but still have not a quantitative metric. As far as we are concerned, it is necessary to present an information-theoretic tool, the balance of information processing (BoIP) to solve this problem.

In this cognitive process, the information flow from two states in time-series, including the redundant, sharing, synergistic information and so on [26]. Nevertheless, computing information flow still include the multiple staggered dependent manner between information categories, but previous studies only divide the information flow into dichotomous components with state-dependent and state-independent in complex tasks [19, 27].To solve this problem, we could consider integrating all information categories as a system-level flow that eliminates redundant information, merges the synergistic information, and balances the interactive information. Instead of segregating it into high coupling components as shown in partial information decomposition (PID) [28]. In fact, the square root of Jensen-Shannon divergence (SRJSD) is a true system-level metric to measure the dissimilarity between two probability distributions [29], which is fully used and explained in our study to compute information flow gain.

As mentioned above, we use and improve the information-theoretic tools, AIS, TE, BoIP to quantify the humans’ state and information processing based on eye-movement. And then, we update a model of system-level information flow between humans and environment, involving those information components whether information categories or relationship have been discovered. The information flow is the pivotal factor to influence humans’ action, prediction, error reduction, and decision-making, even the task performance. [23, 30–32].

## Materials and methods

### Experiment

The present study is designed with a repeated-measures. Each participant completes experiments with the same driving route and the same task instruction, one week apart and a total of 14 *∗* 4 = 56 valid tests, and in a single room without any interference. At the end of the task, they are compensated for their time with CNY 25.

### Participants

We recruit Fourteen Master/Ph.D. students (9 male, 5 females; age range: 21-29, Mean = 21.3, SD = 2.37) at Tianjin University, who participate in the driving experience with a valid driver license for at least one and a half years. All participants report normal or corrected to normal vision and audition, normal color vision, psychiatric health and no history of mental illness. Moreover, none of the participants has adverse reaction to the virtual environment utilized in this study.

### Equipment set-up

Virtual Reality Environment used HTC Vive headset is to display driving scene for participants. The eye-tracking equipment is 7INVENSUN Instrument aGlass DKII [33], embedded into the HTC Vive to capture eye-movement data at a frequency of 90 Hz and an accuracy of gaze position of 0.5°. The driving simulator is Logitech G29 including steering wheel, brake pedals, and accelerator [34].Participants listen to the ambient traffic and car engine sounds in Virtual Environment by speakers. The participants’ visual scanning and driving behaviors are displayed on a desktop monitor (3840 *×* 2160 resolution with the refresh rate at 60 Hz), in which we can observe the experimental procedure. The driving environment is developed by the Unity 3D engine with C SHARP language and uses the UnityCar opensource Plug-in to simulate the car’s structure and power system. And computing raw eye-movement time series data is used 7INVENSUN aGlass SDK for Unity including timestamp, gaze position, hit points position, pupil size. And others such as car speed, position and etc., are used unity SDK.

Furthermore, to ensure high-quality recordings, (i) all participants complete the point calibration procedure prior to the experiments, (ii) the headset is adjusted and fastened to participants’ heads, (iii) sight is adjusted to prevent hair and eyelashes from obscuring, (iv) the seat is adjusted to a comfortable position in front of the simulator and distance within 60-80cm from screens. A participant is experimenting in Fig 1.

**Fig 1.**
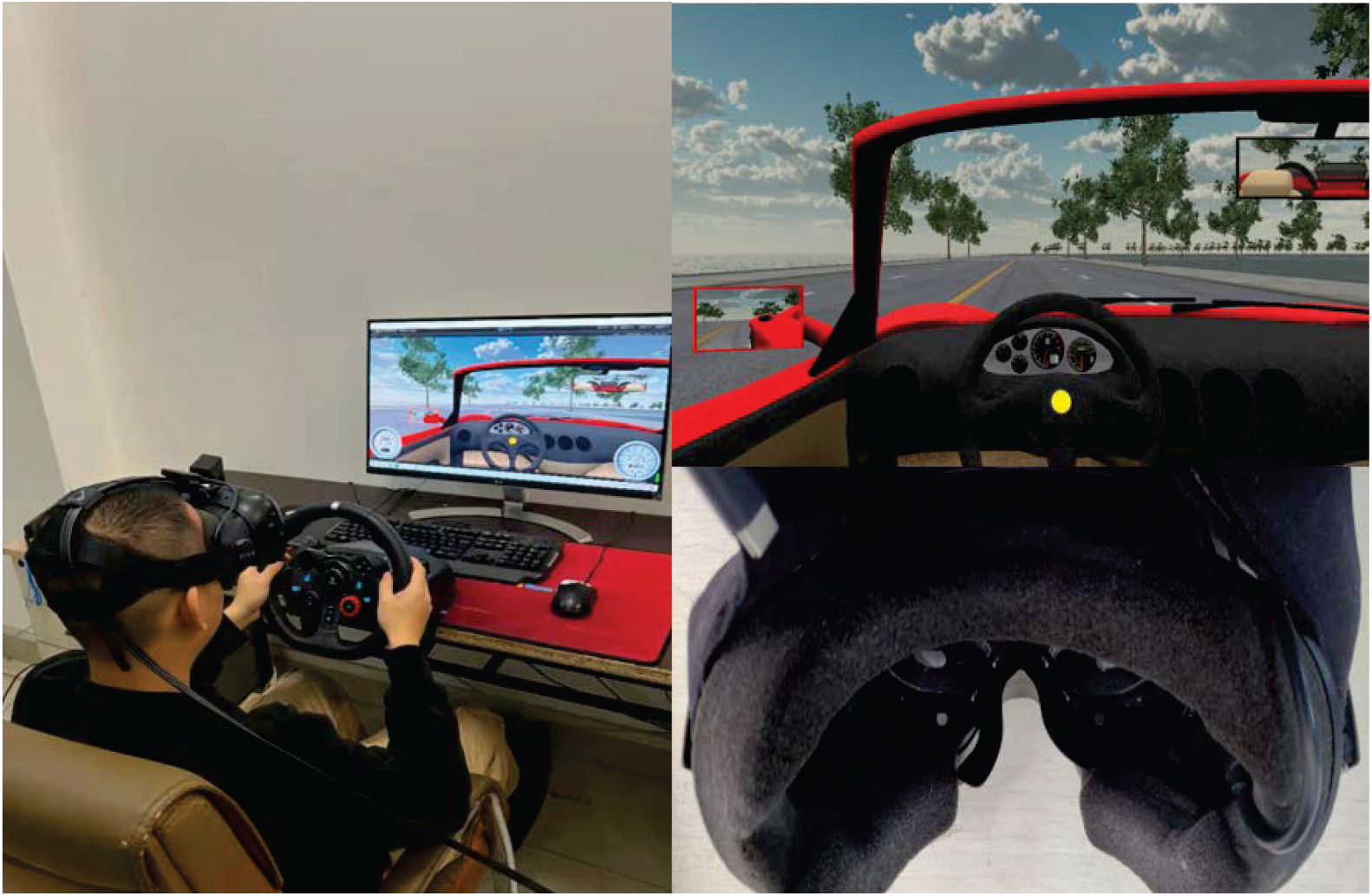
The real and virtual environment and equipment of our experience. Left: real environment. Top right: virtual environment. Bottom right: equipment.

### Procedure in virtual reality environment

Virtual reality experience participants immersed is an effective way to investigate human behavior [35].This study simulates a four-lane two-way suburban road, composed of a succession of nine straight sections and eight bends (4 left bends and 4 right bends with mean radii of curvature ranging from 44 to 313 m), which is realistic and natural without extra variables such as pedestrians, heavy traffic, and other intense stimulator distracting the participants. Similarly, participants only have one goal, to drive the vehicle from starting point to the destination. This goal-directed task helps us better detect general and unblended characterization of cognition and behavior during the driving process [17, 36], such as eye movement, motor action and so on. At the same time, driving-self could be seen as a complex task, which typically happens in everyday life and the driver’s brain continually exchanges information with surroundings as usual [37]. Therefore, it also helps us to explore general driving behavior underlying those characterizations.

Eligible participants are experiment three sessions individually, which are (1) background questionnaire session, (2) training session, (3) driving session. For background questionnaire session, the personal information is collected such as name, age, driving age, and all participants need to complete an Informed Consent Form. For the 10-min training session, participants drive freely without the time and the speed limit to be familiar with the driving simulators, and in which, we observe whether they will be dizzy in the 3D environment to determine whether the experiment could go ahead. After a break of 3–5 min, participants need to check the calibration with the same methodology and adjustment as discussed in section.c, and are instructed that they are going to the next session with the same simulated drive. In this moment, for the 3-min driving session, participants are instructed to comply with driving rules as they do in real life: following the formulated route, limiting the speed at a maximum of 40 km/h, and the lane position close to the center line. In fact, the speed or acceleration of vehicle could be seen as a metric to measure performance for drivers [38].The inverse of mean acceleration as driving performance is used in this paper, so the lower value of mean acceleration, the higher driving performance becomes, and vice versa. Furthermore, recording all data is in this session from beginning to end.

### Fixation and Optical flow

There are different ways to arrive at the destination for each person, similarly different eye-movement strategies to sample environmental information. Primarily, as the one of the patterns of eye-movement, fixation consist of the gaze points with a minimum duration of 100ms and maximum visual angle of 1° dispersion thresholds [39] is the main cognitive unit in decision-making process [40]. The normalized histogram of fixation locations is used to construct the probability distribution of fixations in 3D environment, named fixation distribution, and in this paper, fixation distribution represents the humans (observer) state. In addition, the optical flow resulted from a dynamic environment always happen in driving situations [41], representing a motion variation [42] and the optical flow distribution embedded fixation distribution represents the observation variation on retina (environmental state) [17]. The fixations are calculated using identification by dispersion threshold algorithm(I-DT) [43], and the algorithm of identification of optical flow (see Fig 2 for schematic depiction of fixations and optical flow) and the detail of probability distribution function established are used in [29].

**Fig 2.**
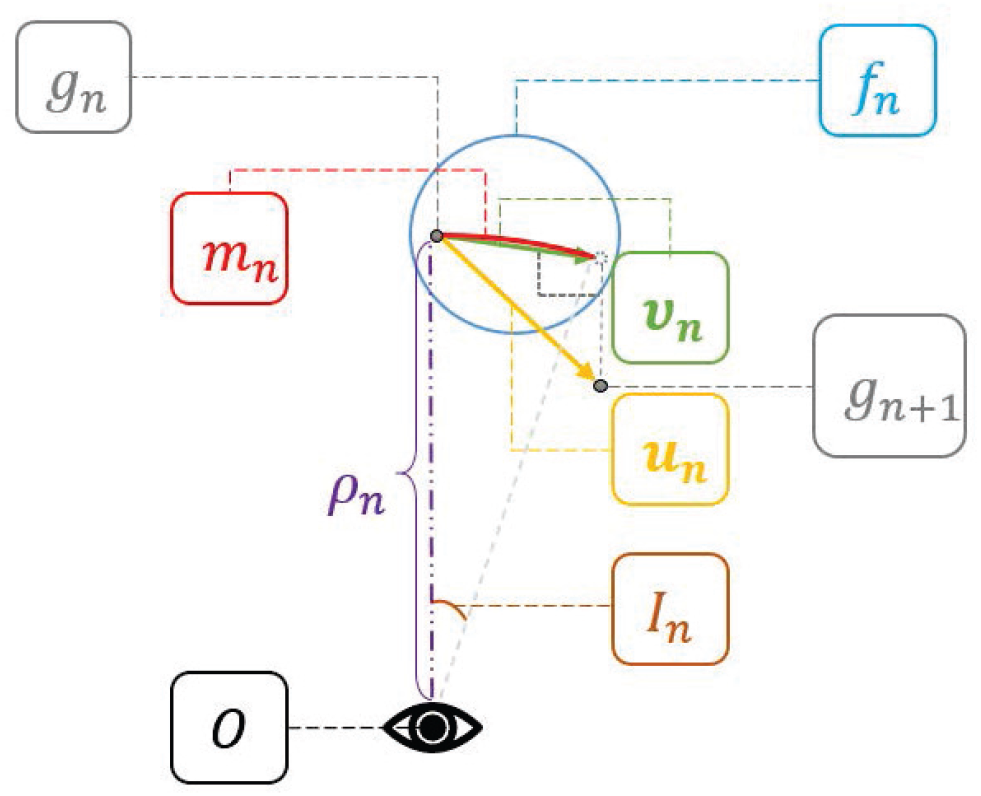
An illustration of optical flow *I*_*n*_ induced by fixation *f*_*n*_.

### Measures of information dynamics

The information dynamics [44] in time series based on Shannon entropy that measures the information quantity by uncertainty reduction, include Transfer Entropy (TE), Active Information Storage (AIS) within complex systems [45], and the Balance of Information Processing (BoIP) we proposed. These measurements include the history information of the time series of the process themselves and the history information of other processes in the system as inputs.

### Active Information Storage and transfer Entropy

Active information storage (AIS) describes that how much information storage in a process *X* from history state *X*(*k*)_t-1_ to target state *X*(*k*)_t_ [44, 46, 47]. In other word, it is defined as the time-directed mutual information in one process [48]:

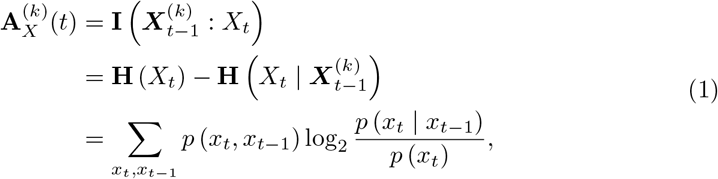

Here *p*(·) and *p*(·|·) denote the (conditional) probabilities of distributions of eye-movement data, and *H*(·), *H*(·|·) and *I*(:) represent the entropy, conditional entropy and the mutual information respectively.Notice that, in *time*(*t*), *X*(*k*)_*t*_ = {*x*_*t−k*_, ⋯, *x*_*t−*1_, *x*_*t*_} are the embedding vectors with history length *k* [49], representing underlying state of process *X* usually for Markov processes. It is clear that a too long history length *k* is served to over-estimate the AIS, so taking the history length as 1 is the common method [50].

Transfer Entropy(TE) formalized by Schreiber [51] and independently by Palus [52], based on conditional mutual information within two subsets {*X*_*t*_, *Y*_*t−*1_ : *t* ∈ *T*} of the individual process in *time*(*t*), is different from AIS. TE quantifying a time-directed transfer of information between two processes by conditioning out shared history information. This definition of TE from source process *Y* to target process *X* also is the case of history length(lag) 1:

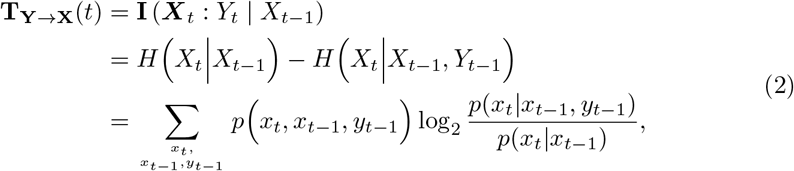

Here, *I*(:| ·) represents conditional mutual information. And analogously this definition also applies to *T*_*X→Y*_ (*t*). *T*_*Y →X*_ (*t*) may be explained intuitively as the uncertainty about *X* determined by past *Y* and *X*, over and above the uncertainty about *X* already determined by its past alone [19]. Note that the values of bidirectional transfer entropy are unequal because the source process, *T*_*Y →X*_ (*t*), provides different history information to the target process. What’s more, the measure of TE may result in a bias of non-zero value. If *T*_*Y →X*_ (*t*) = 0, *X*, conditional on its own past, is independent of the past of *Y*, intuitively, conditional mutual information may be interpreted as a measure of statistical dependence. As a result, we should conduct a statistical significance test to ensure TE not equal to zero. And as shown in Fig 3, provides a sample illustration of AIS and TE.

**Fig 3.**
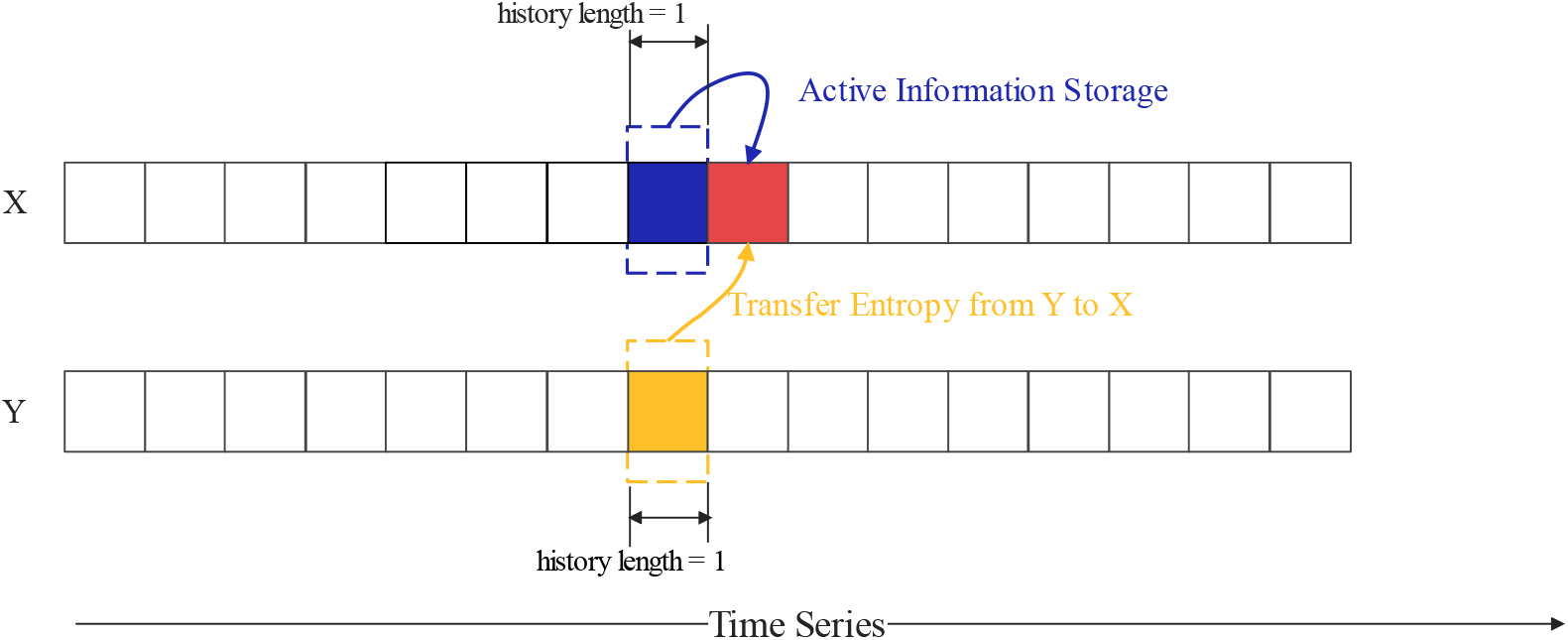
A schematic diagram of the process of active information storage and transfer entropy.

### Balance of information processing

The information will be processed by the humans’ brain, and it is necessary to detect the whose effects after stored and transferred. For the brain, whose flexibility and complexity are the essential properties, which could modulate functional network structure into integrated and segregated brain states when be faced with the demands of the different cognitive challenge tasks. In other words, it is divided into a lattice network (low-activity) and a random structure (high-activity) [45, 53]. Hence, we think that an optimal information processing structure is likely an adjustable balance between the TE and AIS. The transition of order-chaos state is mainly dominated by information storage of the low-activity stage [54] and information transfer of high-activity chaos stage [55, 56]. For driving, the task is complex or high-activity (i.e., the information transfer is dominant). Thus, TE as a denominator, the division of information transfer and storage is proposed:

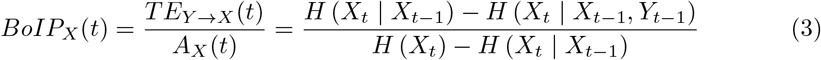

### Measures of system information

It is now well understood that information flow reflects cognitive processes associated with various information, expanding such knowledge to assess behavioral performance in virtual reality settings requires an integrated model of conceptualizing as shown in Fig 4, which is based on the square root of Jensen-Shannon divergence (SRJSD) between two distributions to extract efficiency [29]. The two probability distribution is normalized by studies of [57, 58]. The define of JSD between two distributions *P* = {*p*(*x*)|*x* ∈ *X*} and *Q* = {*q*(*x*)|*x* ∈ *X*} with equal weights [57, 58] is below:

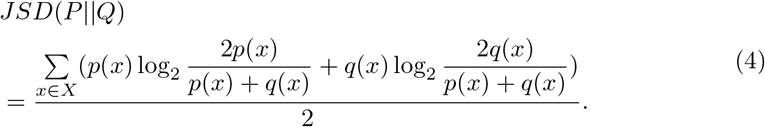

And the true mathematical metric of square root of JSD(P——Q) is used [58].

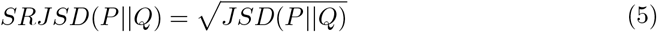

**Fig 4.**
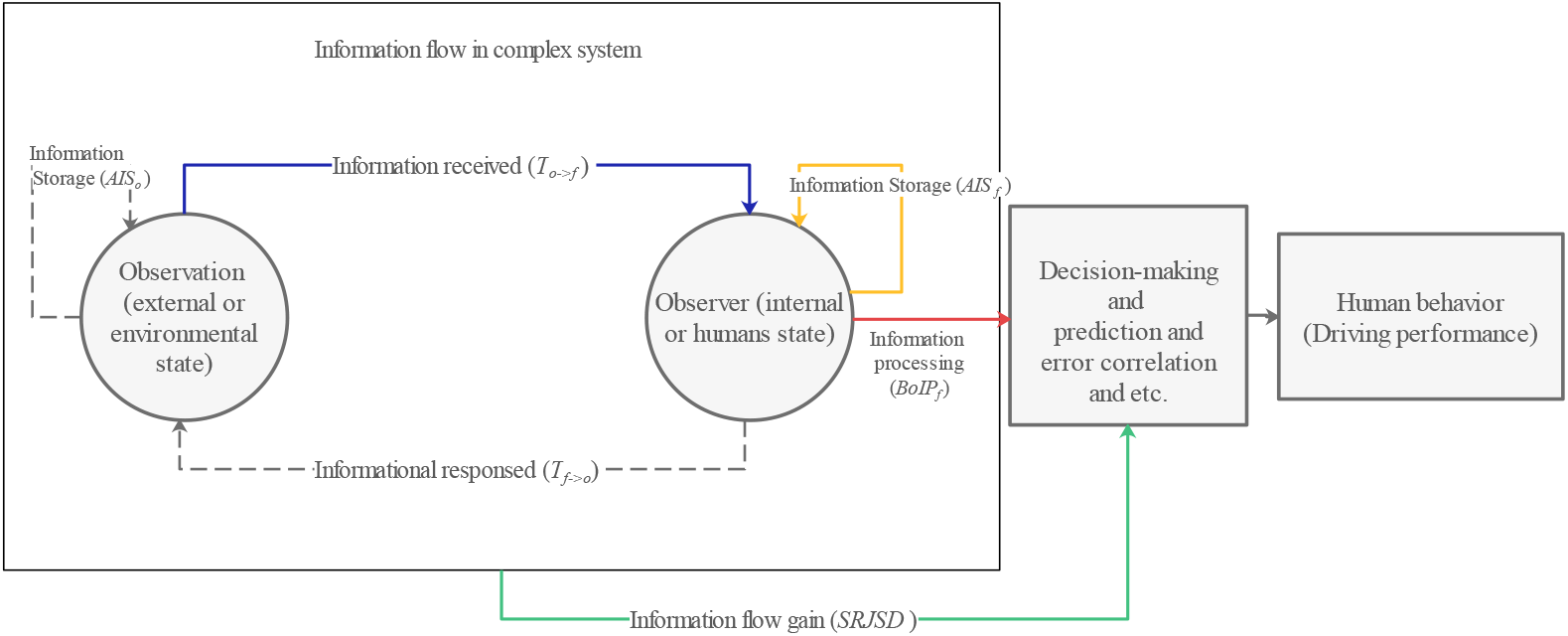
The main process to explain how optical flow and fixation influence human behavior by information flow.

### Significance analysis

In order to assess the statistical significance of the TE and AIS values, it compares to a surrogate distribution of TE and AIS with no direct relationship between the source and the target. The generation of surrogate distribution of TE and AIS is from a number of surrogate source time-series [18, 19]. The generation of surrogate distribution of TE and AIS is from a number of surrogate source time-series. Shuffle the source values to destroy the temporal relationship with the target, and then calculate the corresponding TE and AIS values from each surrogate. The null hypothesis is that there is no direct relationship, we compute a p-value for sampling the actual TE and AIS measurement from the TE and AIS measurements under surrogate distributions by *t − test*. If the test fails, the alternate hypothesis is accepted, which there is a direct relationship. In this study, conducting the test uses 10000 surrogate time-series and a p-value of 0.05 [23].

## Results and discussions

The recent works support that fixation distribution composed of gaze points constrained time and special scope provides an indicator of observers’ state, the same as an internal or human state [9]. For another metric, optical flow distribution, is an external or environmental factor that reflects observation variation due to the motor action [17, 29]. Nevertheless, a problem worth explaining is that the internal and external information produced is not isolated or absolute. These always are regarded as interactive information of two. For a general example in [4], a viewer firstly fixates on a table rather than a bed while searching for coffee cups as whose prior knowledge (internal information). At the same time, the table’s structure, shape, colors, and so on, as external information, help the viewer find the accurate location. So that, the cognitive process goes beyond the standard dichotomy of “bottom-up” (external factors drive the control, such as stimulus salience) and “top-down” (internal factors drive the control to the observer) [25, 59], proved by division of labor of visual cortex, respectively indicating primary visual cortex(V1) encodes bottom-up control settings, V2 encodes top-down control settings, and V4 encodes the interaction between the two.

To sum up, all the information, control setting and modulation mechanism are interactive [25]. Fixation and optical flow distribution combined with information theory, TE, AIS and BoIP may explain the interactive process between humans and the environment. And the symbols of TE, AIS and BoIP between fixation and optical flow distribution established with *A*_*f*_, *A*_*f*_, *T*_*f→o*_, *T*_*o→f*_, and Unless otherwise stated, *BoIP*_*f*_ generally refers to *BoIP*.

### Correlation of AIS with pupil dilation

There are some studies [60–62] have proved that pupil dilation is an indicator of cognitive load or effort. In this paper, we utilize the standard deviation of pupil size during each trial to represent the pupil dilation because of its simplicity and effectiveness [63]. Moreover, according to the information theory, the logarithmic probability of occurrence (*−log*_2_*p*) represents the quantity of information conveyed by the occurrence [35], and then we hypothesize that the *A*_*f*_ may be the central part of cognitive effort. Because the establishment of fixation distribution is correlated with the information that brain continually stored [64]. In this moment, humans will pay more attention [8]. And then the amount of the attentional resource reflects cognitive effort [65].

We used three kinds of widely correlation coefficients (CC): the Pearson linear correlation coefficient (PLCC), Spearman rank-order correlation coefficient (SROCC) and Kendall rank-order correlation coefficient (KROCC), which are employed to conduct correlation analysis. Through these ways, we find the correlation between *A*_*f*_ and pupil dilation (PLCC: *ρ* = -0.41, p ¡ 0.05; SROCC: *ρ* = -0.40, p ¡ 0.05; KROCC: *ρ* = -0.30, p ¡ 0.05), to verify our hypothesis. The negative correlation coefficients mean that the lower information stored in fixation distribution, the higher cognitive effort exerted on humans, which may lead to them seeming confused and nervous. However, this does not mean that the proxy of effort (i.e., pupil dilation or *A*_*f*_) is correlated with task performance directly (correlation between *A*_*f*_ and performance, PLCC: *ρ* = -0.063, p ¿ 0.05; SROCC: *ρ* = -0.097, p ¿ 0.05; KROCC: *ρ* = -0.048, p ¿ 0.05; correlation between pupil dilation and performance, PLCC: *ρ* = 0.158, p ¿ 0.05; SROCC: *ρ* = 0.190, p ¿ 0.05; KROCC: *ρ* = 0.127, p ¿ 0.05). Because the higher effort exerted, the more redundant information may be input, not effective [9]. Especially when drivers feel more nervous, they hope to gather more information to modulate behavior in balance mode. In conclusion, effort is not the directed indicator of task performance but the indirect part, because it still influences the information flow.

### Correlation of TE and BoIP with performance

As mentioned above, the *T*_*o→f*_, *T*_*f→o*_, quantify the interactive information transfer between observation and observer, representing the interactive control setting that will influence the decision-making even task performance [25, 30, 66, 67]. To verify it, we then investigate whether the *T*_*f→o*_ and *T*_*o→f*_, show a correlation with task performance. Indeed, we find the significant correlation between *T*_*o→f*_ and task performance (PLCC: *ρ* = -0.309, p ¡ 0.05; SROCC: *ρ* = -0.330, p ¡ 0.05; KROCC: *ρ* = -0.217, p ¡ 0.05), indicating that the less information transfer into observer, the stronger ability of decision-making even better task performance. The reason for this negative correlation is that the less information that observer receives helps to avoid surprise (self-information) and maintain homeostasis (control its internal state to maintain itself within bounds) [68], so humans are in a more stable state and make more reasonable decisions even perform well. For example, it is easier to keep the original state without surprise for the more experienced driver. The practical information that driver receives is less because they have experimented with so much complex and confusing environment that keep calm naturally. However, for another TE direction, there is no correlation between *T*_*f→o*_ and task performance (PLCC: *ρ* = -0.195, p ¿ 0.05; SROCC: *ρ* = -0.167, p ¿ 0.05; KROCC: *ρ* = -0.108, p ¿ 0.05), we want to detect the reason of this. Firstly, we use the ANOVA to denote the TE between fixation and optical flow distribution that are significant differences (p ¡ 0.01) as illustrated in Fig 5, which suggests that the information received from source states has been changed, because of the different information processing ways for humans and environment. And result also demonstrates that *T*_*o→f*_ is less than *T*_*f→o*_, so the corresponding correlation coefficients is higher and show the significant correlation due to negative correlation. Next, we analyze the connotation of *T*_*f→o*_, the information transferred from fixation distribution to optical distribution is about how observers change to observation variation(i.e., how brain’s information actively transfers into the environment and changes it). One way similar to the example of searching coffee cup [4] is that drivers actively control their gaze move to meaningful scenes in driving environments because of whose sufficient prior knowledge or rich experience. But the participants have requested no mass VR experience in our task. By contrast, they prefer to explore the VR world and be attracted by external environments, so the active control of driver himself/herself is not positive, which may be why *T*_*o→f*_ is work instead of *T*_*f→o*_.

**Fig 5.**
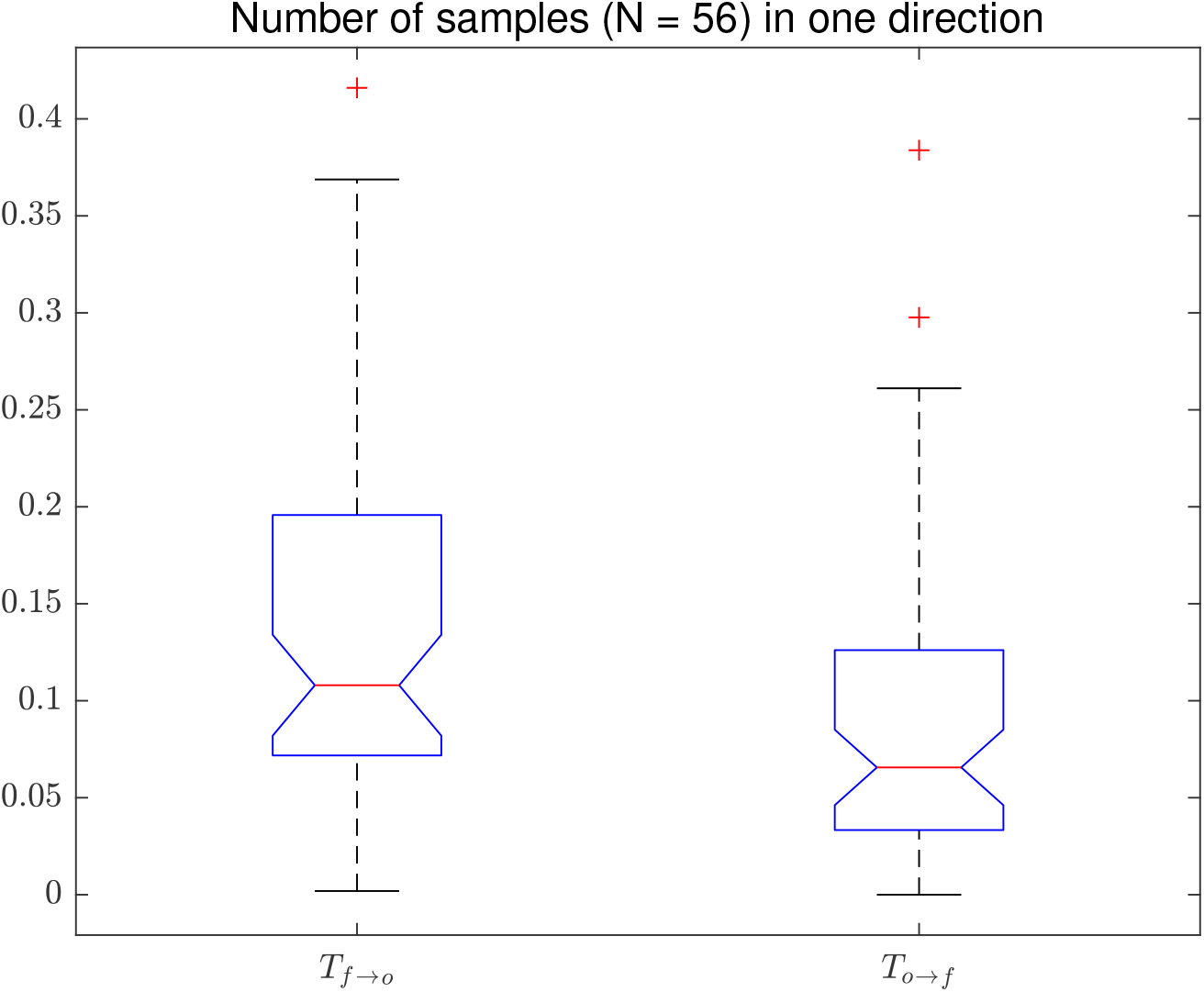
TE between fixation and optical flow distribution. A: From fixation to optical flow. B: From optical to fixation.

For our analysis, not only TE and AIS, the BoIP may correlate with task performance stronger. As mentioned above, the control setting and cognitive effort are the critical roles of balance mode, which is a modulation mechanism that modulate behavior by actively controlling and accepting feedback from cognitive effort passively. Moreover, [24] also studies the relationship between effort and eye-movement, finding that to ensure the more accurate foot-placement that based on effort (cost) and maintain balance, the participants will orient the gaze towards the location that could reduce task-directed uncertainty (i.e., search more information). The balance mode also in [9], implies that an optimal range of modulation extent may influence task performance by selecting the effective samples of visual information through eye-movement. The optimal range means that too strong or too weak modulation will result in worse task performance. In such cases, the ubiquity of high cognitive effort in social anxiety disorder (SAD) and post-traumatic stress disorder (PTSD) due to long-term of focusing on the potential threat (i.e., over-modulation) [69], which has similar aspects with driving in heavy traffic condition. This leads driver to a long-term cognitive load state, too easier distracted even mishandled. But more key information may be ignored when over-relaxed. Thus, there is a balance between control setting and cognitive effort, quantified by BoIP (See 4) that stronger correlation with task performance (PLCC: *ρ* = -0.317, p ¡ 0.05; SROCC: *ρ* = -0.360, p ¡ 0.01; KROCC: *ρ* = -0.248, p ¡ 0.01). Subsequently, as shown in Fig 6, the BoIP curve is smoothed compared to *T*_*o→f*_ and *A*_*f*_ curve, which suggest that the more powerful noise immunity of BoIP, the more stable modulation mechanism that enables better performance.

**Fig 6.**
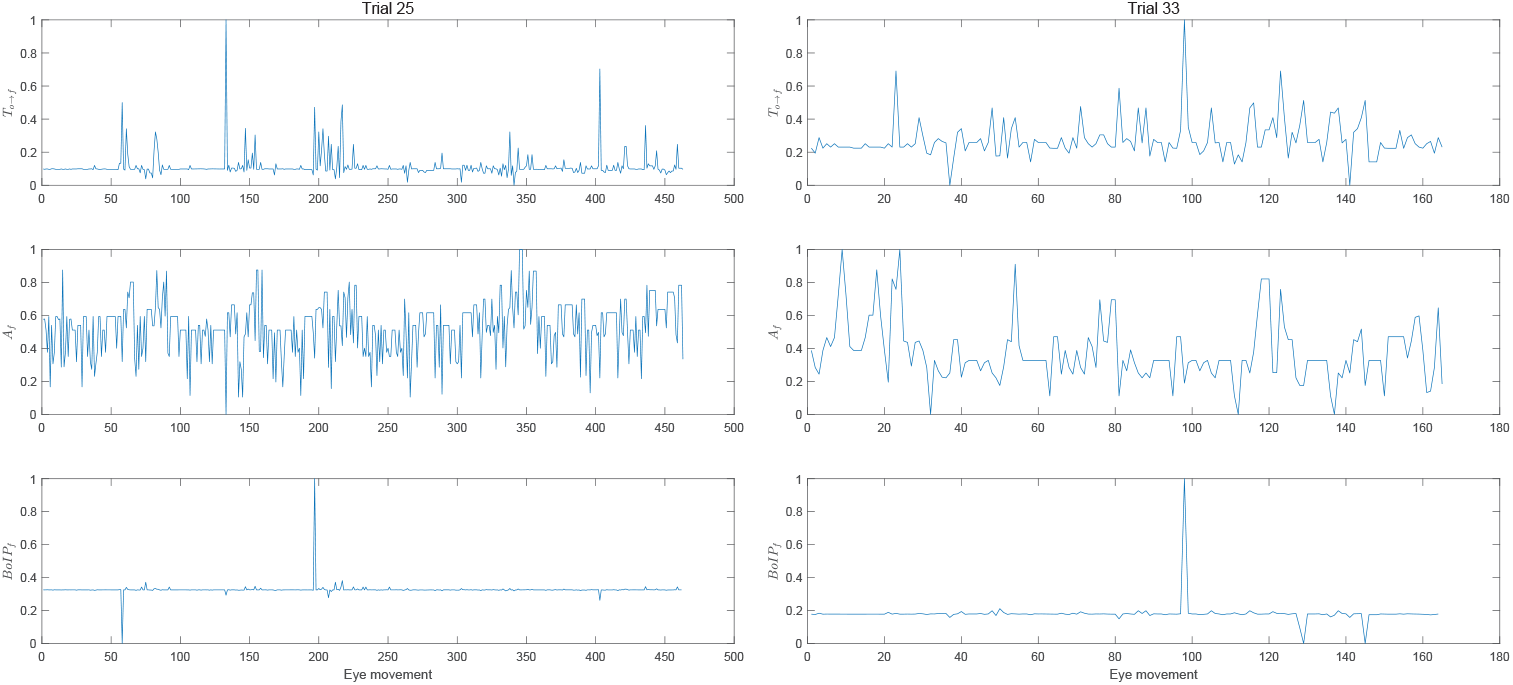
The distribution curves of *T*_*o→f*_, *A*_*f*_ and *BoIP*_*f*_.

### Measurement of information flow gain

The information-theoretic is used to quantify the non-parametric nature and non-linear relationships [52]. Fortunately, the SRJSD is an information metric to quantify system complexity by measuring the dissimilarity between two probability distributions. It is based on Kullback-Leibler divergence as a measurement of information gain [70], with some prominent features that is symmetric or undirected, and identical to the mutual information with a certain condition of [71–73]. As listed in table.1, the SRJSD also shows a statistically significant correlation with task performance. However, in contrast, some direct eye-tracking indices and entropy-based measurements do not. Concretely, GTE [9, 12] is an estimator of uncertainty of scanning pattern. SGE [74] and entropy of fixation sequence (EoFS) [75] consider the uncertainty of spatial pattern. And other fundamental eye-tracking indices based on fixation or saccade, just be appropriate for some particular applications [76]. One essential issue of no correlation between indicators mentioned above and task performance is that not consider the interaction between internal and external factors. Nevertheless, although the TE, AIS, and BoIP especially, have been considered, the CC is a little lower than SRJSD, which is still worth exploring. First and foremost, the framework of information system based on Shannon entropy contains the information storage, information transfer and the balance of two, modelled by AIS, TE and BoIP. Secondly, AIS, TE and BoIP, just represent the unidirectional information flow between observer and observation variation as shown in figure.structure, but the SRJSD quantify the system-level information flow, with some difference that is undirected. Finally, the ultimately effective and remaining information in the system has eliminated redundant and sharing information and merged synergistic information after information processing because of ignoring the multiple staggered dependent manners of information categories. However, AIS, TE, BoIP and other measurements are just a subset of the information flow. Indeed, the information system is explicable precisely if all the subsets could be discovered, but can not do it at present. Thus, this integrated information set is the information flow we understood in complex system not just the concept of information transferred [19]. Besides, the reason for a positive correlation between SRJSD and performance is that the higher SRJSD means that the more efficient information remained in system, which be called information flow gain, representing the stronger ability of humans to modulate behavior [24, 77, 78].

**Table 1.** CC between indicators and driving performance

| Indicators | Pearson |  | Spearman |  | Kendall |  |
| --- | --- | --- | --- | --- | --- | --- |
|  | CC | p-value | CC | p-value | CC | p-value |
| **SRJSD** | **0.49** | **p<0.05** | **0.48** | **p<0.05** | **0.34** | **p<0.05** |
| *BoIP<sub>f</sub>* | -0.34 | p<0.05 | -0.35 | p<0.05 | -0.23 | p<0.05 |
| *T<sub>o→f</sub>* | -0.30 | p<0.05 | -0.33 | p<0.05 | -0.21 | p<0.05 |
| *T<sub>f→o</sub>* | -0.20 | p>0.05 | -0.17 | p>0.05 | -0.11 | p>0.05 |
| GTE | 0.15 | p>0.05 | 0.06 | p>0.05 | 0.04 | p>0.05 |
| SGE | 0.11 | p>0.05 | 0.08 | p>0.05 | 0.05 | p>0.05 |
| EoFS | 0.14 | p>0.05 | 0.12 | p>0.05 | 0.10 | p>0.05 |
| Entropy rate | 0.05 | p>0.05 | 0.08 | p>0.05 | 0.05 | p>0.05 |
| Fixation rate | -0.14 | p>0.05 | -0.06 | p>0.05 | -0.05 | p>0.05 |
| Saccade amplitude | 0.12 | p>0.05 | -0.08 | p>0.05 | -0.05 | p>0.05 |

In conclusion, in the present study, we apply the information-theoretic tools, active information storage (AIS) that quantifies the humans’ cognitive effort, and then propose a metric, balance of information processing (BoIP), combined with AIS and transfer entropy (TE) for the first time to quantify the information within the mode that balances cognitive effort and control setting. These tools and metrics provide a data-driven perspective to understand the information system. And in this study, a major technological innovation is updating a system-level model to analyze humans’ behavior performance, whose effect is more significant than the research on the unique factor by decomposing the information system.

